# Higher synaptic threshold for NMDA spike generation in human neurons

**DOI:** 10.1101/2021.10.22.465509

**Authors:** Guilherme Testa-Silva, Marius Rosier, Suraj Honnuraiah, Robertas Guzulaitis, Ana Morello Megias, Chris French, James King, Katharine Drummond, Lucy M Palmer, Greg J Stuart

## Abstract

Neurons receive synaptic input primarily onto their dendrites. While we know much about the electrical properties of dendrites in rodents, we have only just started to describe their properties in the human brain. Here we investigate the capacity of human neurons to generate NMDA spikes. We find that dendritic iontophoresis of glutamate, as well as local dendritic synaptic stimulation, can evoke NMDA spikes in dendrites of human layer 2/3 pyramidal neurons. Surprisingly, however, the capacity to evoke NMDA spikes in human neurons was significantly reduced compared to that in rodent neurons. Simulations in morphologically realistic models indicated that human neurons have a higher synaptic threshold for NMDA spike generation. Using a simplified model, we show that this is primarily due to the larger diameter of human dendrites. In summary, we find reduced NMDA spike generation in human compared to rodent neurons due to the larger diameter of human dendrites.

## Introduction

Dendrites are the primary site of excitatory and inhibitory synaptic input to neurons in the brain. While dendrites were initially considered passive structures that funnel synaptic signals to the soma and axon (the main output pathway), evidence primarily from rodents indicates that the dendrites of most neurons contain a range of voltage-activated channels that can both actively support and locally generate electrical events (Stuart and Spruston, 2015). A variety of active electrical events have been described in the dendrites of rodent cortical pyramidal neurons, including backpropagating action potentials (Larkum et al., 2007; Stuart and Sakmann, 1994), as well as locally generated dendritic sodium (Larkum et al., 2007; Stuart et al., 1997), calcium (Larkum et al., 1999a; Schiller et al., 1997; Williams and Stuart, 2002) and NMDA (Larkum et al., 2009; Nevian et al., 2007; Palmer et al., 2014; Schiller et al., 2000) spikes. Importantly, these active dendritic properties influence action potential generation in cortical pyramidal neurons (Stuart et al., 1997) and can shift neuronal output from regular to burst firing during distal dendritic excitatory input (Larkum et al., 1999b; Williams and Stuart, 1999). In addition to an impact on action potential output, active dendritic mechanisms in cortical pyramidal neurons are also involved in synaptic plasticity (Gordon et al., 2006; Kampa et al., 2006; Letzkus et al., 2006; Sjostrom and Hausser, 2006) and coincidence detection of proximal and distal synaptic input (Larkum et al., 1999b; Williams and Stuart, 2002). More recent work *in vivo* indicates that active dendritic mechanisms in cortical pyramidal neurons are critical for sensory encoding (Larkum et al., 2009; Palmer et al., 2014; Takahashi et al., 2016), feature detection (Lavzin et al., 2012), orientation tuning during visual stimulation (Smith et al., 2013), correlating sensory and motor input (Xu et al., 2012) and learning (Cichon and Gan, 2015).

In addition to this work in rodents, two recent studies indicated that the dendrites of human cortical pyramidal neurons are also electrically excitable and can show both action potential backpropagation and locally generated sodium/calcium spikes (Beaulieu-Laroche et al., 2018; Gidon et al., 2020). What is not known is whether the dendrites of human neurons also generate NMDA spikes, which are important for local synaptic integration in thin dendritic processes such as basal and apical tuft dendrites (Larkum et al., 2009; Nevian et al., 2007; Schiller et al., 2000). This is the focus of the current study, where we investigate the capacity of human neurons to generate NMDA spikes and compare this with rodent neurons.

## Results

Whole-cell current-clamp recordings were performed from the soma of layer 2/3 (L2/3) pyramidal neurons in brains slices prepared from human tissue. Human brain tissue was obtained fresh from patients undergoing neurosurgery to remove brain tumors or treat epilepsy (performed by Drs. Drummond and King, respectively). This tissue was either excess to that required for histopathology or removed incidentally for surgical access to the brain tumour. Written informed consent was obtained from patients (aged 18 to 70 years old), and all procedures were performed with the approval of the Human Research Ethical Committee (HREC) of the Royal Melbourne Hospital (HREC No: 2001:85 and 2020:214). Human brain tissue was obtained primarily from the anterior medial temporal cortex, as well as the frontal and parieto-occipital cortex was macroscopically healthy and normal and was processed immediately on removal. Consistent with this, human L2/3 pyramidal neurons had resting membrane potentials around −65 mV (average: −67.7 ± 0.5 mV, n=39), fired repetitive action potentials and had morphology typical of cortical L2/3 pyramidal neurons (Figure 1A; Supplemental Figure 1).

**Figure 1:**
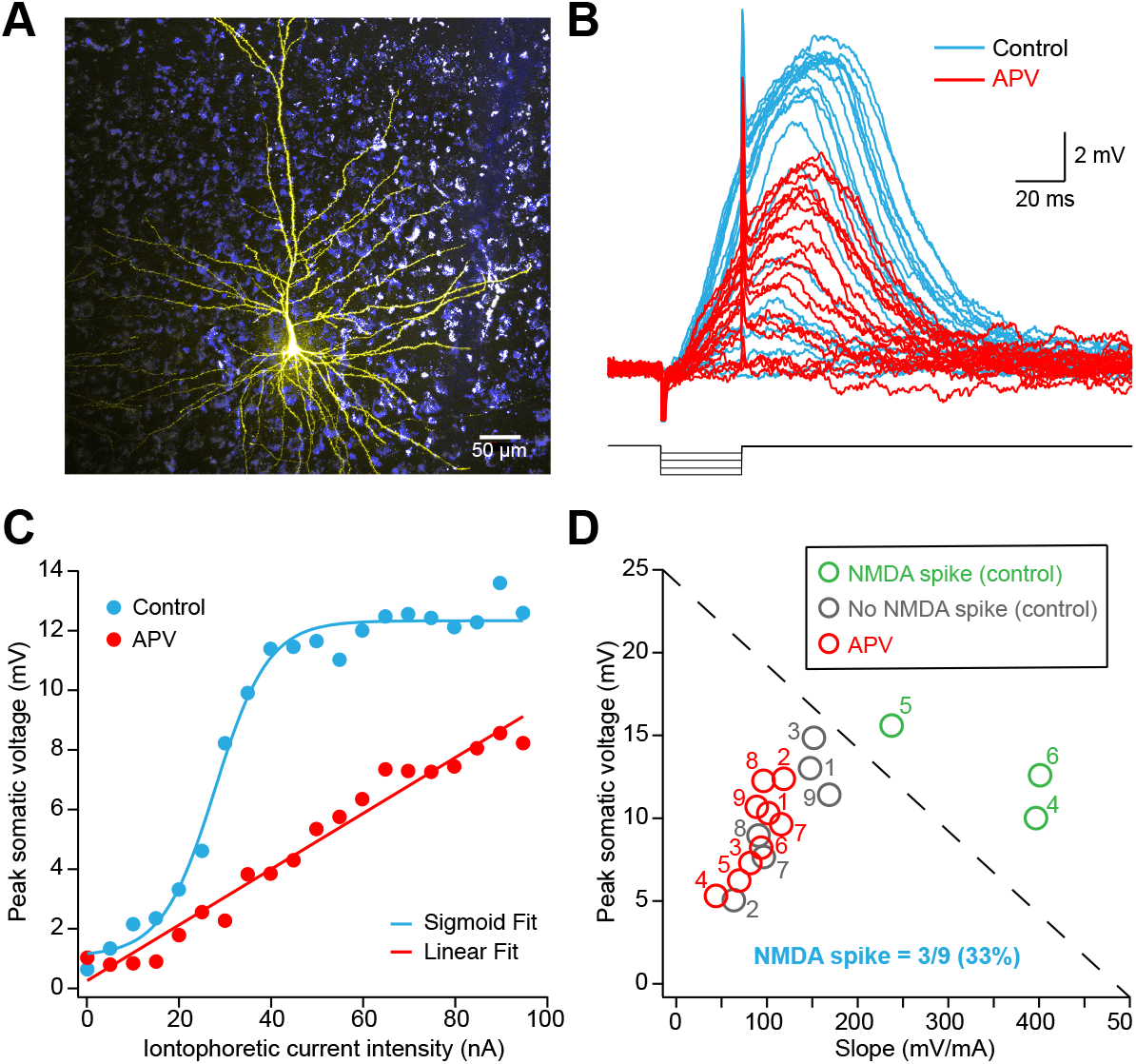
NMDA spike generation in human L2/3 cortical pyramidal neurons. (A) Image of a human L2/3 cortical pyramidal neuron filled with fluorescent dye via the somatic whole-cell recording pipette. (B) Somatic voltage response during glutamate iontophoresis to a basal dendrite (∼150 μm from the soma; 0 to 100 pA in 5 pA steps) in a human L2/3 cortical pyramidal neuron in control (blue) and following bath application of the NMDA antagonist APV (red; 50 μM). The timing and amplitude of the iontophoretic current is indicated schematically (bottom). (C) Plot of the peak somatic voltage at the soma versus iontophoretic current intensity for a human L2/3 cortical pyramidal neuron in control (blue) and APV (50 μM, red). Same cell as shown in panel B. Application of APV (a NMDA antagonist) converted the sigmoidal voltage versus iontophoretic current relationship in control to a linear relationship. Control data fitted with a sigmoid. Data in APV fitted with a line. (D) Plot of the maximum somatic voltage response at the highest iontophoretic current intensity versus the maximum slope of the sigmoid fit in control or the slope of the linear fit in APV for human L2/3 cortical pyramidal neurons (n=9). The dashed diagonal line represents a linear “threshold” used to differentiate cells that had a clear non-linear component under control conditions (green; indicating generation of an NMDA spike) from cells where the response was linear under control conditions (grey) and in APV (red). Individual cells before and after APV are numbered. Only 3 out of 9 (33%) human L2/3 cortical pyramidal neurons showed clear NMDA spikes during dendritic glutamate iontophoresis.

To investigate if human L2/3 pyramidal neurons generate dendritic NMDA spikes we focally activated dendritic NMDA receptors using iontophoresis. Briefly, a pipette containing the neurotransmitter glutamate was placed under visual control in close proximity (within a few μm) to basal and oblique dendrites and brief (20 or 30 ms) negative current pulses of different magnitudes were used to apply different amounts of glutamate locally. We have previously used this method to reliably evoke NMDA spikes in the basal dendrites of rodent cortical pyramidal neurons (Bock and Stuart, 2016). Only cells where we could obtain data before and after application of the NMDA receptor blocker APV were included for analysis (n=9 out of 39 human neurons). This technique reliably depolarized both basal and oblique dendrites (Figure 1B). Importantly, in some cases the amplitude of depolarizing events was non-linearly related to the magnitude of iontophoretic current used to apply glutamate, suggesting the generation of NMDA spikes (Figure 1B,C, blue). Consistent with this idea, bath application of APV (50 μM) converted this non-linear relationship into a linear relationship (Figure 1B,C, red). Control data were fitted with a sigmoid, whereas data in the presence of APV was fitted with a line. From these fits, we determined the maximum slope of the sigmoid in control as well as the slope of the line in APV (see Methods). For each cell we then plotted the sigmoidal slope in control versus the maximum voltage obtained during iontophoretic glutamate application (Figure 1D). A linear “threshold” was then used to differentiate cells that had a non-linear component under control conditions (Figure 1D; green) from cells where the response under control conditions was essentially linear (Figure 1D; grey). The rationale was that cells that show a clear non-linearity, indicative of the generation of an NMDA spike, would be expected to have a high sigmoidal slope in control and high maximum amplitude, so will be located in the upper righthand quadrant. In contrast, cells where NMDA spikes were not evoked under control conditions would be expected to have reduced sigmoidal slope and to reduced maximum amplitude, so will be located in the bottom left-hand quadrant. As evidence of the validity of this approach, the maximum sigmoidal slope and maximum amplitude in control were significantly reduced by application of APV in cells that showed an NMDA spike, whereas they should be similar in cells that did not show an NMDA spike (Figure 1D; red). Based on this analysis, we conclude that only 3 out of 9 (33%) human L2/3 pyramidal neurons showed a clear NMDA spike during dendritic glutamate iontophoresis. These data, for the first time, indicate that human neurons can generate NMDA spikes.

Since most of our current knowledge about dendritic NMDA spikes stems from work in rodents during synaptic input, we directly compared the capacity of human and mouse L2/3 pyramidal neurons to generate NMDA spikes evoked by local dendritic extracellular stimulation. As the majority of the human data was from temporal cortex, mouse recordings were also made from temporal cortex. Only cells where we could obtain data before and after application of the NMDA receptor blocker APV were included for analysis (n=8 out of 18 human neurons). These experiments revealed that NMDA spikes could be evoked by extracellular stimulation of basal dendrites in both human (Figure 2A) and rodent (Figure 2B) neurons. Consistent with this idea, somatic responses to local dendritic extracellular stimulation increased in a sigmoidal manner with stimulation intensity under control conditions in some cells, whereas the relationship between extracellular stimulation strength and response amplitude were linear in the presence of the NMDA antagonist APV (Figure 2C,D). We applied the same “threshold” analysis used to classify cells as having an NMDA spike or not during iontophoretic glutamate application to this extracellular stimulation data (see Methods). This analysis differentiated human and mouse neurons that had a clear non-linear component during extracellular stimulation under control conditions (Figure 2E,F; green), indicative of generation of an NMDA spike, from cells where the response was essentially linear (Figure 2E,F; grey). Similar to the experiments during glutamate iontophoresis, only a small subset of human L2/3 cortical pyramidal neurons showed clear NMDA spikes during local dendritic extracellular stimulation (Figure 2E; 2 out of 8; 25%). In contrast, the majority of mouse L2/3 cortical pyramidal neurons showed clear NMDA spikes (Figure 2F; 11 out of 15; 73%). These data illustrate that the capacity to evoke NMDA spikes in human L2/3 cortical pyramidal neurons is considerably lower than in mice L2/3 cortical pyramidal neurons.

**Figure 2:**
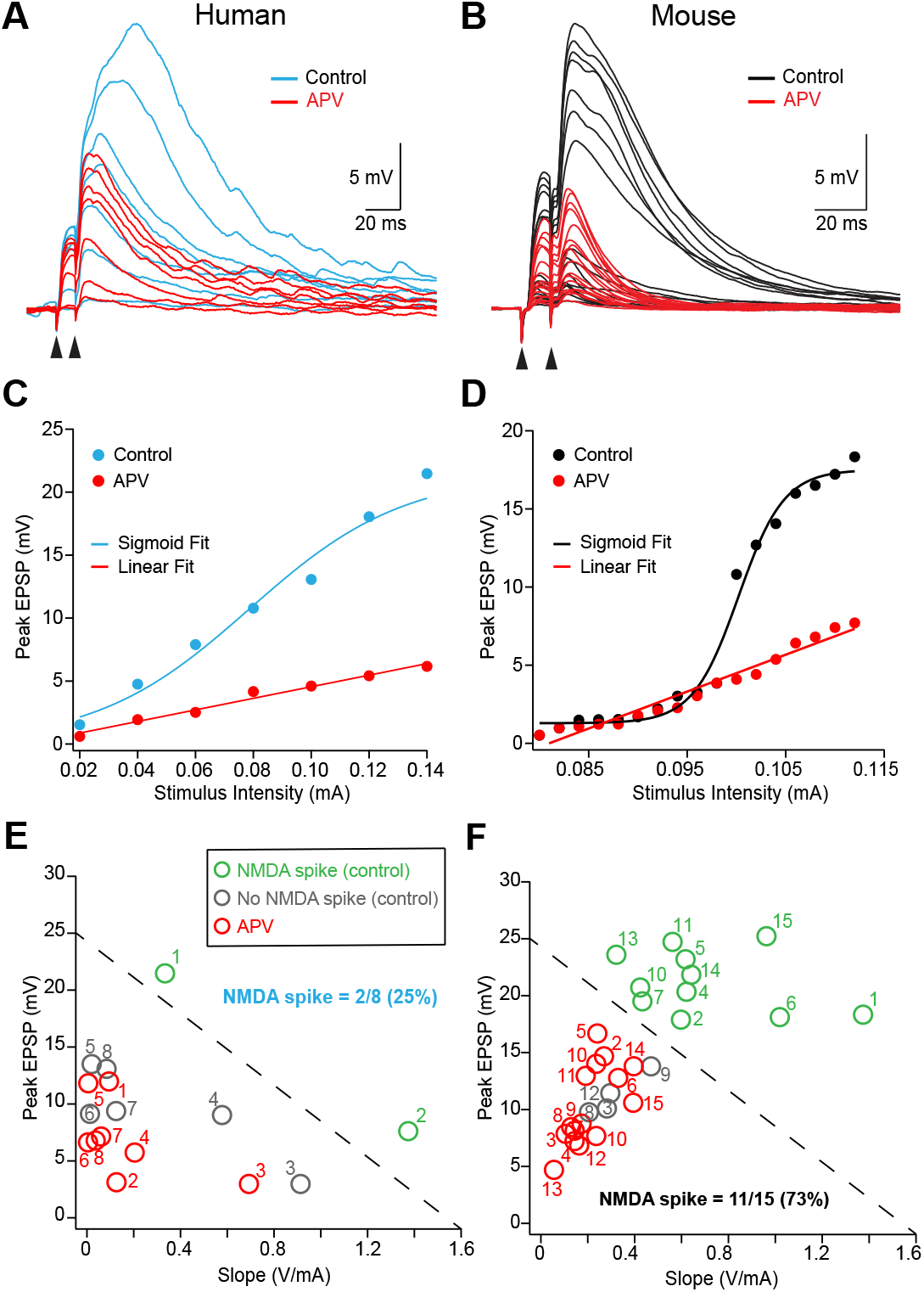
Comparison of NMDA spike generation in human versus mouse L2/3 cortical pyramidal neurons. (A,B) Examples of voltage responses at the soma during local extracellular stimulation (two pluses at 100 Hz) to basal dendrites of human (A) and mouse (B) L2/3 cortical pyramidal neurons at different intensities in control (blue/black) and APV (50 μM, red). (C,D) Plot of the peak somatic voltage response versus extracellular stimulation current for human (C) and mouse (D) L2/3 cortical pyramidal neurons in control (blue/back) and APV (50 μM, red). Application of APV (an NMDA antagonist) converted the sigmoidal somatic voltage response versus stimulus current relationship in control to a linear relationship. Control data fitted with a sigmoid. Data in APV fitted with a line. Human data from the cell shown in panel A. Mouse data from the cell shown in panel B. (E,F) Plot of the maximum somatic voltage response at the highest extracellular stimulation intensity versus the maximum slope of the sigmoid fit in control or the slope of the linear fit in APV for human (E, n=8) and mouse (F, n=15) L2/3 cortical pyramidal neurons. The dashed diagonal line represents a linear “threshold” used to differentiate cells that had a clear non-linear component under control conditions (green; indicating generation of an NMDA spike) from cells where the response was linear under control conditions (grey) and in APV (red). Individual cells before and after APV are numbered. Only 2 out of 8 (25%) human L2/3 cortical pyramidal neurons showed clear NMDA spikes (E). In contrast, 11 out of 15 (73%) mouse L2/3 cortical pyramidal neurons showed NMDA spikes (F).

We next examined dendritic integration in morphologically realistic models of human and mouse L2/3 cortical pyramidal neurons (Figure 3A; see Methods). Different numbers of synaptic inputs containing AMPA and NMDA components were randomly distributed over a defined region of the basal dendrites (10 μm in length, 160 μm from the soma). Intracellular resistance (R_i_) and specific membrane capacitance (C_m_) were set to the same value, with the specific membrane resistance (R_m_) in the human and mouse models adjusted so that the somatic input resistance in the models was similar to that observed experimentally (Supplemental Figure 1; see Methods). Apart from the difference in R_m_, the length of human basal and apical dendrites were clearly longer than those in mouse cortical L2/3 pyramidal neurons (Figure 3A), consistent with earlier work (Mohan et al., 2015). In addition, we found that the diameter of basal dendrites in human L2/3 cortical pyramidal models had a significantly larger diameter (Figure 3B; p < 0.001; Unpaired t-test).

**Figure 3:**
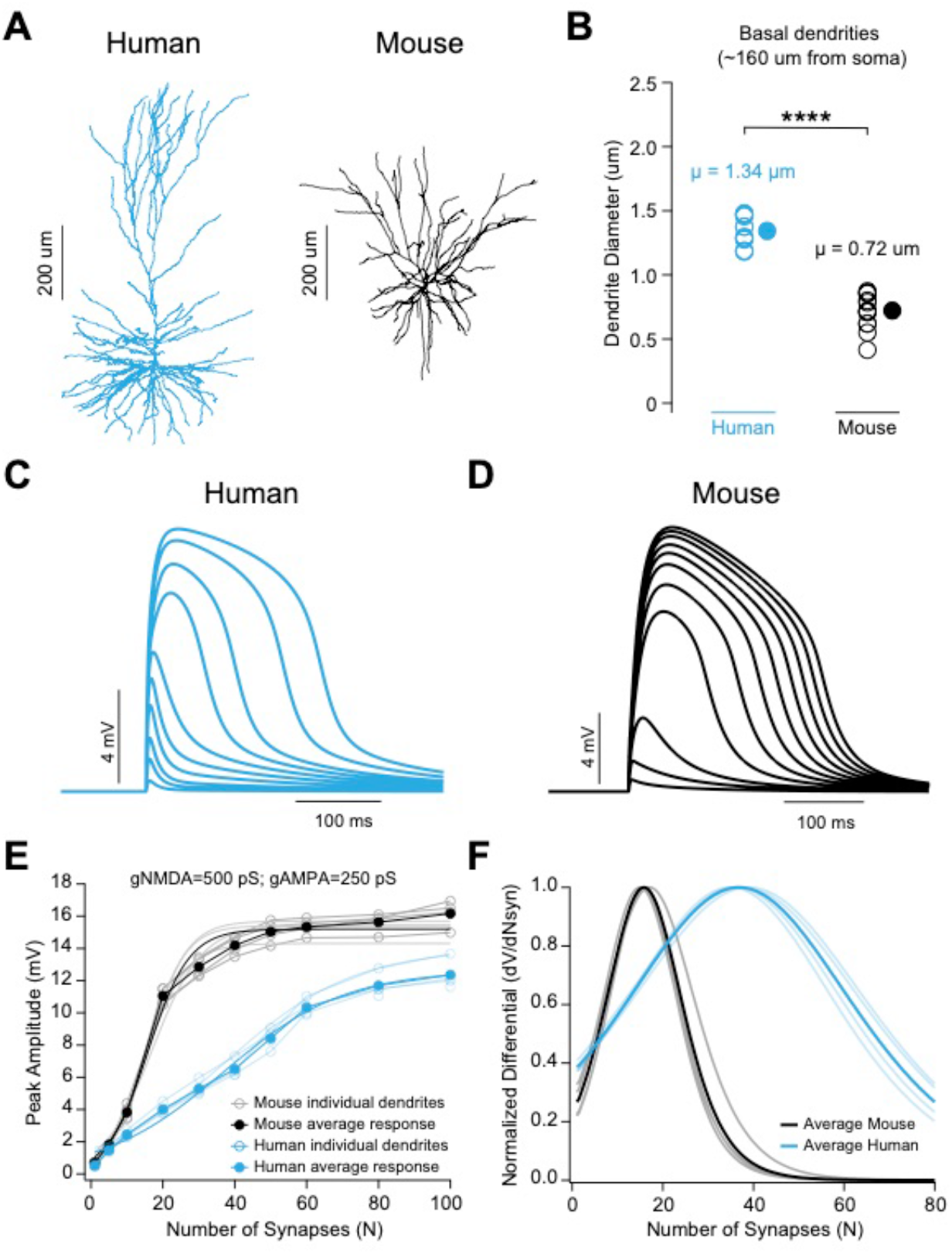
NMDA spike generation in morphologically realistic models of human and mouse L2/3 cortical pyramidal neurons. (A) Images of the human (left) and mouse (right) L2/3 pyramidal neuron models. (B) Comparison on the individual (open symbols) and average (closed symbol) basal dendritic diameter ∼160 μm from the soma in the human and mouse L2/3 pyramidal neuron models. (C,D) Voltage responses at the soma during activation of different numbers of synaptic inputs to basal dendrites in a 10 μm length of dendrite 160 μm from the soma in the human (C) and mouse (D) L2/3 cortical pyramidal neuron models shown in panel (A). (E) Plots of the peak somatic voltage responses versus the number of activated synapses (N) in the human (blue) and mouse (black) L2/3 cortical pyramidal neuron models shown in panel (A) during synaptic input onto different basal dendrites. Light lines with open symbols show responses in individual basal dendrites. Darker lines and filled symbols show the average. All data fitted with sigmoid functions. (F) Plots of normalized, differentiated sigmoidal fits for the data shown in (E) versus the number of activated synapses (N) in the human (blue) and mouse (black) L2/3 cortical pyramidal neuron models shown in panel (A) during synaptic input onto different basal dendrites. Light lines show responses in individual basal dendrites. Darker lines show the average.

Could these differences account for the reduced capacity to generate NMDA spikes in human L2/3 cortical pyramidal neurons? To investigate this, we activated increasing numbers of randomly activated synaptic inputs on different basal dendrites until an NMDA spike was evoked in the human (Figure 3C) and mouse (Figure 3D) L2/3 cortical pyramidal neuron models. The peak amplitude of the response at the soma for different basal dendrites was plotted versus the number of activated synapses and the data fitted with a sigmoidal function (Figure 3E). To determine the number of synaptic inputs required to evoke an NMDA spike we differentiated the sigmoidal fits (Figure 3F). We defined the “threshold” number of synaptic inputs required for NMDA spike generation as the number at the peak of the differentiated sigmoid, which represents its midpoint. This analysis revealed that the number of synapses required to evoke NMDA spikes in the basal dendrites of human L2/3 cortical pyramidal neurons was significantly higher than in mouse basal dendrites (Figure 3F). This observation was consistent across a range of different basal dendrite morphologies in the human and mouse L2/3 cortical pyramidal neuron models (Figure 3E,F). These simulations in morphologically realistic models align with our experimental data and suggest that the morphology of human L2/3 cortical pyramidal neurons, together with their passive properties, are sufficient to explain the reduced capacity to evoke NMDA spikes in human compared to mouse L2/3 cortical pyramidal neurons.

We next used a simplified “ball and stick” model to investigate which passive and morphological parameters underlie the difference in NMDA spike generation in human and mouse L2/3 cortical pyramidal neurons. The model consisted of a somatic compartment connected to a single dendritic compartment. Synaptic input containing both AMPA and NMDA components was placed 160 μm from the soma. The strength of synaptic input was adjusted by changing the magnitude of the combined AMPA/NMDA conductance. The dimensions of the dendritic compartment or its passive membrane properties were then varied over a physiologically relevant range to determine which parameters had the greatest impact on the threshold for NMDA spike generation. The peak amplitude of the evoked response in the somatic compartment was measured during increases in synaptic conductance in models with different dendritic diameters (Figure 4A), dendritic lengths (Figure 4B), R_m_ (Figure 4C), C_m_ (Figure 4D) and spine density (Figure 4E). Different spine densities were modelled by decreasing R_m_ and increasing C_m_ in the dendritic compartment by a scaling factor (Holmes, 1989; Holmes and Rall, 1992). For each model we fitted a sigmoid to the relationship between the amplitude of the somatic response and the magnitude of the synaptic conductance (Figure 4, left) and then differentiated the sigmoid fit (Figure 4, middle). As in Figure 3, we determined the synaptic “threshold” for NMDA spike generation based on the peak of the differentiated sigmoid in the different models. Finally, we plotted the threshold synaptic conductance for NMDA spike generation versus the different passive and morphological parameters examined to determine which had the greatest impact (Figure 4, right). These simulations indicated that differences in dendritic diameter have the greatest impact on NMDA spike threshold (Figure 4A), with differences in dendritic length (Figure 4B), membrane capacitance (Figure 4D) and spine density (Figure 4E) having a much smaller impact. Differences in R_m_ over the range used in the morphologically realistic models had almost no impact on NMDA spike threshold (Figure 4C). Increasing dendritic diameter from 0.75 um to 1.25 um, which is similar to the observed difference in basal dendritic diameter in the mouse and human L2/3 cortical pyramidal neuron models (Figure 3B), was sufficient to double the synaptic threshold for NMDA spike generation in the ball and stick model (Figure 4A). This difference in NMDA spike generation is similar to that seen in the full model (compare Figure 4A to Figure 3F). These simulations therefore indicate that differences in basal dendritic diameter alone can explain the difference in NMDA spike threshold observed in the human and mouse morphologically realistic neuronal models. In summary, using realistic and simplified models we show that the reduced capacity to generate NMDA spikes in human compared to mouse L2/3 cortical pyramidal neurons is likely to primarily be a direct result of the wider diameter of basal dendrites in human neurons.

**Figure 4:**
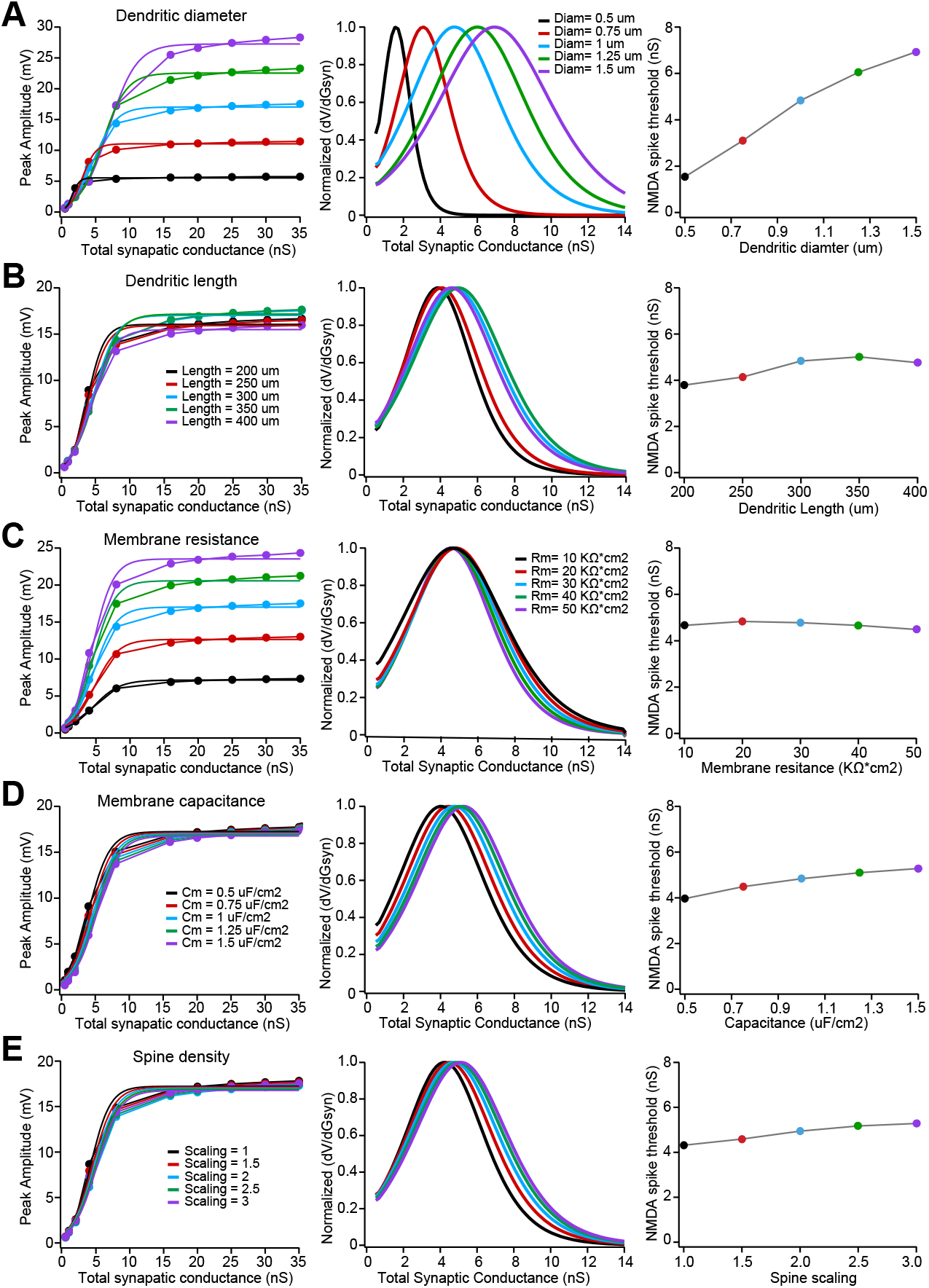
Impact of different passive and morphological parameters on NMDA spike generation in a simplified “ball and stick” model. (A – E) Left, Plots of the peak voltage response in the somatic compartment together versus the total AMPA/NMDA synaptic conductance for models with different dendritic diameter (A), dendritic length (B), specific membrane resistance (C; R_m_), specific membrane capacitance (D; C_m_) and spine density (E). All data fitted with a sigmoid. Middle, Plots of the differentiated and normalized sigmoid fits show on the left versus the total AMPA/NMDA synaptic conductance for models with different dendritic diameter (A), dendritic length (B), specific membrane resistance (C; R_m_), specific membrane capacitance (D; C_m_) and spine density (E). Right, Plots of NMDA spike threshold versus dendritic diameter (A), dendritic length (B), specific membrane resistance (C), specific membrane capacitance (D) and the spine density scaling factor (E). NMDA spike threshold was defined as the AMPA/NMDA synaptic conductance at the peak of the differentiated sigmoid fits (middle). Spine density was adjusted by dividing R_m_ and multiplying C_m_ by the indicated scaling factor. Data color coded as indicated in the legends.

## Discussion

Understanding how neurons in the human brain process information is critical to an understanding of how the human brain works. Here we show that similar to previous work in rodents, the dendrites of human cortical pyramidal neurons can generate NMDA spikes. Surprisingly, however, the ability to evoke NMDA spikes in human L2/3 cortical pyramidal neurons was significantly reduced compared to that in mouse neurons. This finding was reproduced in morphologically realistic models, where the number of synaptic inputs required for NMDA spike generation was significantly larger in the human L2/3 pyramidal neuron model. Using a simplified model, we show that the synaptic threshold for NMDA spike generation is highly sensitive to differences in dendritic diameter. Together, these simulations suggest that the reduced capacity to generate NMDA spikes in human compared to mouse cortical L2/3 pyramidal neurons results from the larger dendritic diameter of human basal dendrites.

Quantitative analysis of the morphology of human and rodent cortical pyramidal neurons indicates that basal dendrites of human neurons are significantly longer than those in rodents (Mohan et al., 2015). While this may contribute to the higher synaptic threshold for NMDA spike generation in human neurons, our simulations indicate this effect was small (Figure 4B). In contrast, lower C_m_ values, which have been reported in human neurons (Eyal et al., 2016), acted to decrease rather than increase NMDA spike threshold, although this effect was also small over the physiological range (Figure 4D). Furthermore, more recent work suggests that C_m_ in human neurons is similar to that in rodents (Beaulieu-Laroche et al., 2018). As expected from these simulations, where dendritic spines were accounted for by increasing C_m_ while lowering R_m,_ changing C_m_ increased the threshold for NMDA spike generation. Human cortical L2/3 pyramidal neurons are thought to have a higher spine density than mouse L2/3 pyramidal neurons, with human spines larger than mouse spines (Benavides-Piccione et al., 2002). This leads to a doubling of the surface area of basal dendrites in human cortical pyramidal neurons (Deitcher et al., 2017), equivalent to a spine scaling factor of 2 in our simulations. While increasing the density of dendritic spines increased the threshold for NMDA spike generation, as with the impact of changing dendritic length this effect was small (Figure 4E). In contrast, changes in dendritic diameter had by far the greatest impact on the threshold for NMDA spike generation, which was essentially doubled by an increase in dendritic diameter from 0.75 to 1.25 μm (Figure 4A). This difference in NMDA spike threshold was similar to that seen in human and mouse morphologically realistic models (Figure 3F). We conclude from this that the primary factor underlying differences in NMDA spike generation in human and mouse neurons is the observed difference in basal dendrite diameter (Figure 3B). The idea that basal dendrites of human L2/3 pyramidal neurons have larger diameter than in the mouse is also supported by the finding that they have a higher spine density (Benavides-Piccione et al., 2002) given previous work indicating that spine density is linearly related to dendritic diameter (Larkman, 1991).

Increasing basal dendritic diameter increased the amplitude of NMDA spikes in the somatic compartment of the ball and stick model (Figure 4A, left). In contrast, the amplitude of NMDA spikes in human and mouse L2/3 pyramidal neurons was similar (Figure 2E,F; Human 14.9 ± 2.0 mV versus mouse 17.5 ± 1.6 mV; P = 0.352; Unpaired t-test), as was the case in the human and mouse morphologically realistic models (Figure 3C,D). What underlies this apparent discrepancy? The most likely explanation is that differences in charge transfer (which underlie the dependence of NMDA spike amplitude on dendritic diameter in the ball and stick model) are offset by differences in input resistance in real neurons. While the ball and stick simulations with different dendritic diameter have similar input resistance, the input resistance of human L2/3 pyramidal neurons (35.9 ± 2.3 MΩ, n = 39) was significantly lower than that of mouse L2/3 pyramidal neurons (88.36 ± 2.64 MΩ; n = 59; p < 0.001; Wilcoxon-Mann-Whitney rank test). As a result, reduced charge transfer in thinner mouse dendrites is offset by their higher input resistance leading to a similar NMDA spike amplitude to that seen in human neurons.

Recent work indicates that, like rodent neurons, the dendrites of human cortical pyramidal neurons support active action potential backpropagation as well as the generation of local sodium/calcium spikes (Beaulieu-Laroche et al., 2018; Gidon et al., 2020). Our findings complement this earlier work by showing that human neurons can also generate local dendritic NMDA spikes, due to regenerative activation of NMDA receptors. In contrast to work in rodents, however, the earlier work by Beaulieu-Laroche and colleagues (Beaulieu-Laroche et al., 2018) found that dendritic spikes generated by distal apical dendritic current injection in human cortical layer 5 (L5) pyramidal neurons were weaker and remained more compartmentalized compared to locally generated dendritic spikes in rodents. This was attributed to the increased dendritic length of human cortical L5 pyramidal neurons compared to rodents and a reduced density of dendritic voltage-activated channels. Consistent with these findings, the work by Gidon and colleagues (Gidon et al., 2020) found that action potential backpropagation in human cortical L2/3 pyramidal neurons was weaker than in rodent neurons. Furthermore, dendritic calcium spikes generated by dendritic current injection were brief and graded in amplitude. These observations concur with our finding that NMDA spike generation is reduced in human compared to mouse cortical pyramidal neurons. While one can never rule out that factors such as differences in tissue quality, the long-term impact of medication or the older age of the human tissue may explain these differences with rodent neurons, our findings suggest that human dendrites are less excitable than rodent dendrites in part simply due to their larger size.

## Methods

Human brain tissue excess to that required for histopathology or removed incidentally for surgical access was obtained from patients undergoing neurosurgery to remove brain tumors or treat epilepsy (performed by KD and JK, respectively). Written informed consent was obtained from patients (aged ∼20 to 70 years old) and all procedures were performed with the approval of the Human Research Ethical Committee (HREC) of the Royal Melbourne Hospital (HREC No: 2001:85 and 2020:214). The tissue obtained is from the anterior medial temporal cortex, the frontal cortex and the parieto-occipital cortex. After resection, tissue was placed immediately (< 30s) in a cold and recently carbogenated solution containing (in mM): 125 NaCl, 25 NaHCO_3_, 5 HEPES, 1 CaCl_2_, 6 MgCl_2_, 3 KCl, 1.25 NaH_2_PO_4_ and 10 Glucose. Samples were transported on ice until the slicing procedure, which occurred generally less than 15 minutes after resection. Rodent tissue was obtained from 4-6 weeks old mice. Mice were anesthetised by inhalation of isoflurane (3%) and decapitated prior to the brain being removed according to procedures approved by the Animal Ethics Committee of the Australian National University.

Brain tissue was sliced using a vibrating tissue slicer (Leica Microsystems) in ice-cold carbogenated solution containing (in mM): 110 Choline Chloride, 26 NaHCO_3_, 11.6 Na-Ascorbate, 7 MgCl_2_, 3.1 Na-pyruvate, 2.5 KCl, 1.25 NaH_2_PO_4_, 0.5 CaCl_2_ and 10 Glucose. Coronal issue slices (300 to 400 μm thick) were obtained containing temporal, frontal or parieto-occipital cortex. Slices were incubated in carbogenated solution containing (in mM): 125 NaCl, 25 NaHCO_3_, 5 HEPES, 1 CaCl_2_, 6 MgCl_2_, 3 KCl, 1.25 NaH_2_PO_4_ and 10 Glucose at 35 °C for 30 minutes and thereafter at room temperature until required. Recordings were performed in artificial cerebral spinal fluid (ACSF) with composition (in mM): 125 NaCl, 25 NaHCO_3_, 3 KCl, 1.2 CaCl_2_, 0.7 MgCl_2_, 1.25 mM NaH2PO4 and 10 Glucose.

### Electrophysiology

Whole-cell patch-clamp recordings were made from the soma of L2/3 pyramidal neurons using an Olympus BX51 WI microscope equipped with a fluorescent imaging system. Slices were continuously perfused with carbogenated ACSF at a rate of 2ml/min at 34°C (±1°C). Borosilicate glass pipettes (inner diameter 0.86 mm, outer diameter 1.5 mm, Sutter) were pulled using an electrode puller (Sutter Instruments, USA) and had open tip resistances of 4 - 6 MΩ. Recording pipettes were filled with intracellular solution of the following composition (in mM): 130 K-Gluconate, 10 KCl, 10 HEPES, 4 Mg^2+^-ATP, 0.3 Na_2-_GTP, 10 Na_2_-Phosphocreatine (pH set to 7.25 with KOH, osmolarity 285 mosmol/l). 5 μM of the fluorescent dye Alexa 594 or 0.02% TMR biocytin (Thermo Fisher Scientific) was included in the intracellular solution to visualise basal dendrites. Current clamp recordings were made using a current-clamp amplifier (BVC-700A; Dagan Corp., Minneapolis, MN or Multiclamp 700B; Molecular Devices, San Jose, USA) mounted on a remotely controlled micromanipulators (Luigs and Neumann, Germany). Cells were excluded from data analysis if the somatic resting membrane potential was more depolarised than −55 mV or if fluctuations in membrane potential > 5 mV were observed at any time during the recording. Cells were also excluded if the somatic series resistance exceeded 50 MΩ or if the somatic series resistance changed by more than 10% during the recording.

### Glutamate iontophoresis

L-Glutamic acid (Tocris, UK) was dissolved in ACSF to a concentration of 200 mM and the pH adjusted to 7.4 with NaOH. 1 μM of the fluorescent dye Alexa 594 was added to the glutamate solution to help visualise the pipette tip under fluorescent light. Iontophoretic pipettes were made from borosilicate glass (outer diameter of 1 mm; inner diameter of 0.5 mm) and pulled to give a tip resistance of 120-140 MΩ when filled. The iontophoretic pipette was connected to the 1x gain headstage of an Axoclamp 2A current/voltage-clamp amplifier (Molecular Devices, Sunnyvale, CA). A backing current of −5 nA was used to prevent spillage of glutamate from the iontophoretic pipette. The pipette was inserted into the slice and positioned under visual control in close proximity (within 5 μm) of a basal or oblique dendrite of the patched neuron at a location at least 120 μm from the soma (average ∼160 μm from the soma). Fluorescence was visualised using a CCD camera (CoolSNAP EZ, Photometrics, Tucson, AZ). Glutamate were applied using 20 to 30 ms negative current steps of varying amplitude.

### Extracellular stimulation

Extracellular stimulation pipettes were pulled from theta glass (outer diameter 1.5 mm, Sutter) to approximate tip diameters of 1 to 2 μm. Both barrels were filled with carbogenated ACSF containing 5 μM of the fluorescent dye Alexa 594 and connected to the positive and negative poles of an isolated stimulator (WPI, Getting Instruments or ISO-Flex). The pipette tip was positioned under visual control in close proximity (within 2 μm) of a basal dendrite of the patched neuron at a location at least 120 μm from the soma (average ∼160 μm from the soma). Brief (0.2 ms) pulses of different amplitude were used to evoke synaptic responses.

### Data acquisition and analysis

Electrophysiological data were filtered at 10 kHz and acquired at 50 kHz by a Macintosh computer running Axograph X acquisition software (Axograph Scientific, Sydney, Australia) using an ITC-18 interface (Instrutech/HEKA, Germany), or by a Windows computer running Clampex 10.7 (Molecular Devices) using a Digidata 1440A interface (Molecular Devices, San Jose, USA). Data analysis was performed using Axograph X in combination with IGOR Pro, or using Clampfit 10.7. Pooled data are presented as mean ± standard error of the mean (SEM).

Plots of EPSP/NMDA spike amplitude versus iontophoretic current or extracellular stimulation intensity in control were fitted with a sigmoidal relationship, whereas data in the presence of APV was fitted with a line. The maximum slope of the sigmoid in control was determined by differentiating the sigmoidal fit and finding its peak. For each cell we then plotted the maximum slope of the sigmoid in control or the slope of the linear fit in APV versus the maximum voltage obtained during the highest iontophoretic current or greatest extracellular stimulation intensity used (see plots in Figure 1D and Figure 2E,F). The peak of differentiated sigmoidal fits was also used to determine the “threshold” number of synaptic inputs or synaptic conductance required to evoke an NMDA spike in the models.

### Modelling

Computer simulations were performed using the NEURON 7.4 simulation environment (Carnevale and Hines, 2006). The morphology of the human neuron was obtained from Eyal et al. (Eyal et al., 2018), whereas the mouse L2/3 pyramidal neuron was obtained from Smith et al. (Smith et al., 2013). C_m_ was set to 1 μF/cm^2^ and internal resistance (R_i_) 150 Ω.cm. R_m_ was set to 10,000 Ω.cm^2^ in the human and 40,000 Ω.cm^2^ in the mouse L2/3 pyramidal neuron model, giving somatic input resistances of 40 MΩ and 100 MΩ, respectively. These values are similar to that observed experimentally (human L2/3 neurons: 35.9 ± 2.3 MΩ, n = 39; mouse L2/3 neurons: 88.36 ± 2.64 MΩ; n = 59). The resting membrane potential was set to −65 mV. Dendritic compartments were not adjusted for spines and did not contain voltage-activated channels. Different numbers of synaptic inputs containing both AMPA (250 pS) and NMDA (500 pS) components were randomly placed on basal dendrites within a restricted dendritic region (10 um in length) with its center 160 μm from the soma. Simulations used an integration time constant of 25 μs.

In the simplified passive model, a somatic compartment (dimensions: 90 μm by 90 μm) was connected to a single dendritic compartment with default dimension of 300 μm length and 1 μm width. The default passive properties were R_m_ = 30,000 Ω.cm^2^, C_m_ = 1 μF/cm^2^ and R_i_ = 150 Ω.cm. The resting membrane potential was set to −65 mV. A single synaptic input of varying magnitude with an AMPA to NMDA conductance ratio of 2 was placed in the dendritic compartment 160 μm from the somatic compartment. In different simulations the dimensions of the dendritic compartment, as well as R_m_ and C_m_, were varied systematically to investigate how these parameters impact on the magnitude of the synaptic conductance required to generate an NMDA spike. Different spine densities were modelled by decreasing R_m_ and increasing C_m_ in the dendritic compartment by a scaling factor (Holmes, 1989; Holmes and Rall, 1992). Simulations used an integration time constant of 25 μs.

## Acknowledgements

This research was funded by an Australian Research Council Discovery Project grant to GJS and LMP (DP190103296) and the Sylvia and Charles Viertel Charitable Foundation (LMP).

## Author Contributions

GJS, LMP and GTS initiated and designed the project. GTS, MR, SH, AMM & RG acquired and analyzed the experimental data. JK & KD obtained the human tissue. CF coordinated tissue collection and provided experimental expertise. SH performed the modeling. GJS & LMP supervised the project. GJS & SH prepared the figures and GJS, LMP, MR, GTS, SH and RG wrote the paper, which was edited by all authors.

