## Supplementary figures and images for "Higher synaptic threshold for NMDA spike generation in human neurons"

### Supplemental Figure 1

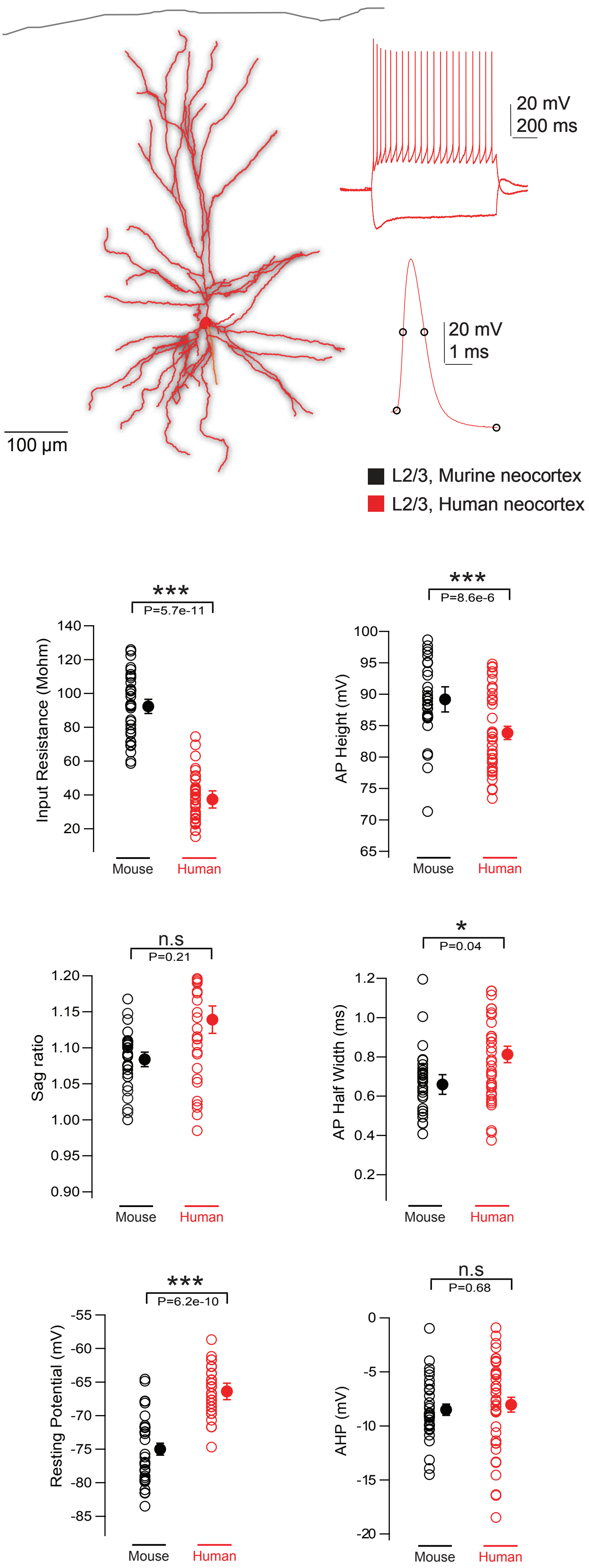
